# Hierarchical Gating of Cortical Population Dynamics Drives Pain

**DOI:** 10.64898/2026.03.06.710170

**Authors:** Eric Hu, Guanghao Sun, Elaine Zhu, Chengju Tian, Isabel Reyes, Zhe Sage Chen, Arjun V. Masurkar, Qiaosheng Zhang, Jing Wang

## Abstract

The prefrontal cortex and anterior cingulate cortex are key cortical hubs for pain regulation, yet the functional hierarchy between them remains unclear. We examined how the prelimbic cortex (PL) to ACC pathway regulates nociceptive processing and pain behavior in freely moving rats across synaptic, cellular, and network levels. Activation of the PL to ACC pathway reduced aversion to both evoked and spontaneous pain, whereas inhibition increased pain aversion, indicating that this circuit exerts tonic modulatory control. Meanwhile, ex vivo electrophysiology revealed that PL inputs recruit local ACC interneurons to produce feedforward inhibition of ACC pyramidal neurons. At the cellular level, in vivo microendoscopic calcium imaging showed that optogenetic activation of PL axon terminals suppresses nociceptive-evoked activity of ACC pyramidal neurons. At the network level, PL activation reduced pain-induced excitability while centralizing nociceptive information flow within the ACC, resulting in a gated, low output population state. Overall, these findings identify a hierarchically organized cortical circuit that tonically controls pain related sensory and affective experience.

## Introduction

Physiological responses to sensory inputs require coordinated integration of activity across multiple cortical regions. The prefrontal cortex (PFC) and anterior cingulate cortex (ACC) are central regions in this process, playing key roles in a wide range of sensory and affective behaviors, including depression, addiction, and pain, across species^1,2^. However, despite their importance, surprisingly little is known about how these two regions are hierarchically organized in regulating complex behaviors.

Studies in both humans and animal models have identified the ACC as a critical neural substrate for the affective – or aversive – experience of pain^3–13^. ACC neurons increase their firing rates in response to nociceptive inputs^14–20^, and the neural activity in this region has been shown to decode the intensity and timing of pain^12,21–25^. In rodent models, inhibiting or lesioning the ACC has resulted in decreased aversion to noxious stimuli in conditioned place aversion (CPA) assays, whereas activation of this region is associated with enhanced pain aversion ^8,12,13,26,27^.

Like ACC neurons, individual neurons in the PFC are recruited into specific functional neural circuits in a context-dependent manner^28^. Prefrontal nociceptive processing involves multiple neurotransmitter systems, including glutamatergic, endocannabinoid, and cholinergic signaling^29–32^, and extensive work has mapped efferent projections to subcortical areas such as the periaqueductal gray and nucleus accumbens^33–38^. In rodents, the prelimbic cortex (PL) shares functional homology with human dorsolateral prefrontal cortex, which is implicated in both acute and chronic pain modulation^39–41^. Like ACC neurons, PL neurons show increased firing rates in response to noxious stimuli^42^. In contrast to ACC neurons, activated PL pyramidal neurons primarily project to subcortical structures to inhibit, rather than facilitate, pain behaviors^36,37,42^.

Thus, the current literature indicates that while neurons in both regions respond to nociceptive inputs, the ACC has a facilitating, whereas the PL has an inhibitory role. Anatomic projections from the PL to the ACC are well-documented^43^, raising the question of whether the PL, beyond its well-known subcortical projections, could also regulate pain by modulating the function of ACC neurons. In this study, we dissected this cortico-cortical projection in the regulation of pain aversion. Activation of presynaptic inputs from PL neurons suppressed ACC population responses to nociceptive stimuli, through a reduction in a global decrease in single neuron activity, coupled with centralization of nociceptive information flow within the ACC. At the synaptic level, this suppression can be mediated via recruitment of local inhibitory circuits. Importantly, optogenetic activation of the PL to ACC projection reduced aversion to both acute and persistent pain, whereas inactivation of this circuit enhanced it, indicating a tonic modulatory role. Together, these findings reveal a hierarchical organization in which the PFC gates pain – and potentially other sensory–affective processes – through top-down control over other cortical circuits.

## Results

### Activation of the PL-ACC projection reduces aversion to both evoked and spontaneous pain

Given the well-documented role of ACC activity in regulating the affective pain experience^12,13,23,46^, we tested whether this projection from the PL to the ACC could modulate pain-aversive behaviors. We injected anterograde CaMKII-ChR2 vs. eYFP control into the left PL, followed by optical fiber implantation in the right ACC, allowing us to specifically activate the presynaptic terminals of PL neurons in the ACC (Fig. 1A). We then performed a series of conditioned placepreference (CPP) tests, which have been used extensively to assess aversive pain responses^12,23,37,44,47,48^. In a two-chamber device, rats received 20 minutes of place-dependent conditioning with either noxious PP, or PP combined with blue light activation of the PL-ACC projection, followed by a 10-minute testing period where neither PP nor optogenetic stimulation was delivered (Fig. 1B). Time spentin each chamber was calculated using video tracking software. After conditioning, the ChR2 cohort of rats preferred the chamber associated with optogenetic activation of the PL-ACC projection (Figs. 1C-D), suggesting that the activity of this pathway effectively reduces the aversive response to acute pain. In contrast, we found no place preference within the eYFP control cohort (Figs. 1E-F).

**Figure 1:**
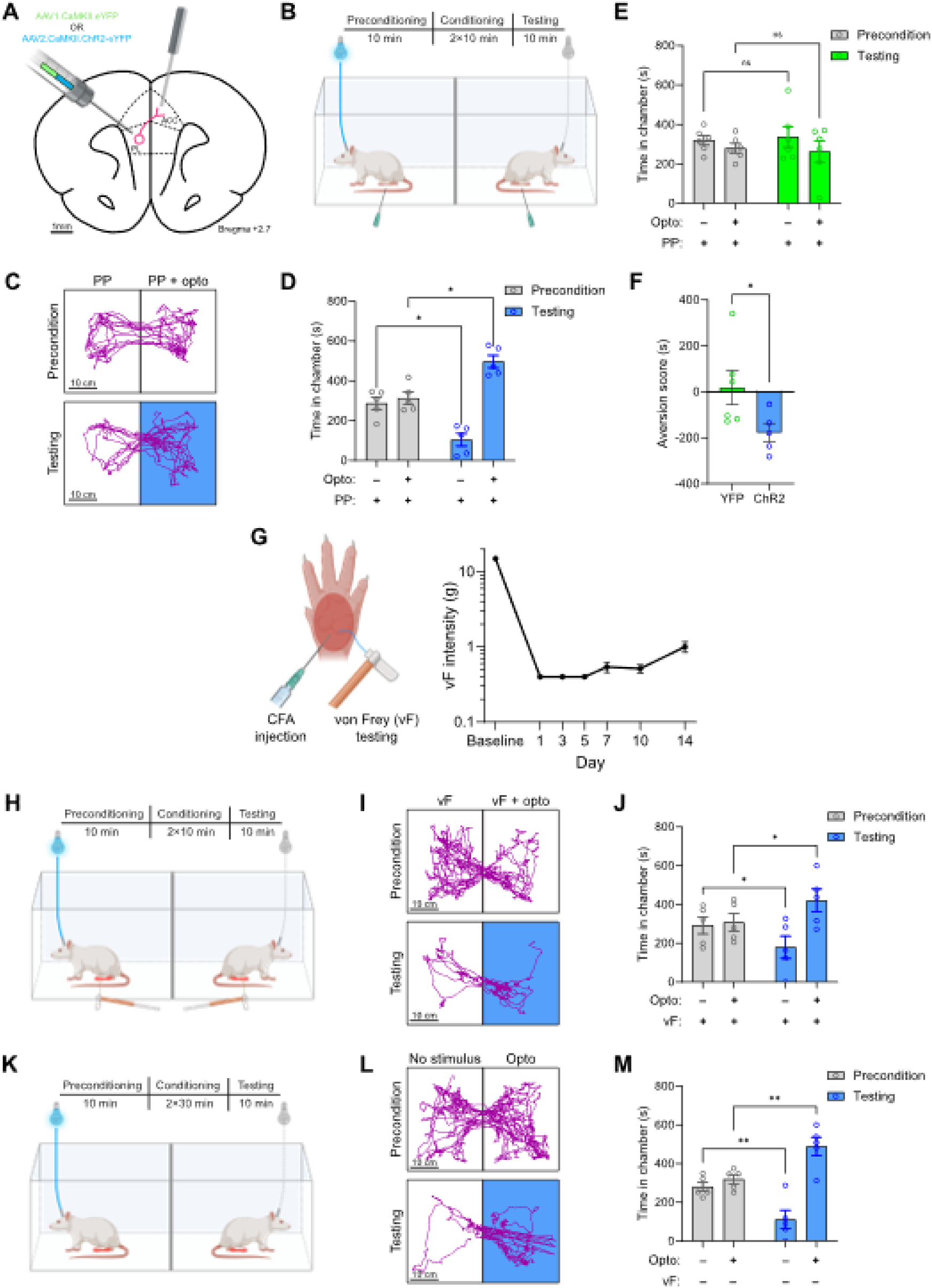
PL-ACC activation reduces pain-aversive behavior. **A** Control/excitatory viral injection protocol and optical fiber implantation. **B** CPP setup and experimental protocol for PP-paired conditions in acute pain model. **C-D** In the presence of acute PP, subjects demonstrate preference for the chamber associated with optoactivation of PL-ACC (panel **C**: representative locomotion traces; panel **D**: p = 0.0135, n = 5, paired t-test). **E** Optoactivation in YFP-injected subjects has no effect on pain aversion (p = 0.820, n = 6, paired t-test). **F** ChR2 PL-ACC optoactivation decreases aversion scores compared to YFP controls (p = 0.0496, n = 5 vs 6, unpaired t-test w/ Welch’s correction). **G** (**Left**) Diagram of footpad CFA injection and von Frey (vF) monofilament stimulus delivery. (**Right**) 50% paw withdrawal thresholds in the mechanical allodynia test indicate strong hypersensitivity to ordinarily non-noxious von Frey monofilament up to 14 days post CFA injection (n = 5, p < 0.0001, one-way ANOVA with Dunnett’s multiple comparisons test). **H** CPP setup and experimental protocol for 6g vF-paired conditions in chronic pain model. **I-J** In the presence of acute vF, chronic pain subjects demonstrate preference for the chamber associated with optoactivation of PL-ACC (panel **I**: representative locomotion traces; panel **J**: p = 0.043, n = 5, paired t-test). **K** CPP setup and experimental protocol for no-stimulus conditions in chronic pain model. **L-M** In the absence of acutely noxious stimuli, chronic pain subjects demonstrate preference for the chamber associated with optoactivation of PL-ACC (panel **L**: representative locomotion traces; panel **M**: p = 0.0093, n = 5, paired t-test).

To test if this pathway also regulates persistent or chronic pain, we applied a well-known inflammatory model by injecting complete Freund’sadjuvant (CFA) into the left hindpaw^12,49^ (Fig. 1G). Mechanical allodynia testing was conducted over the course of two weeks after CFA injection, to confirm that subjects indeed developed hypersensitivity (or allodynia) to ordinarily non-noxious stimulus with von Frey monofilament (vF) (Fig. 1G), followed by CPP experiments, were during this hypersensitive window (9 days after CFA). We used a 6g vF to evoke allodynia in both treatment chambers, while pairing one chamber with optogenetic activation of the PL-ACC projection, and the opposite chamber with no activation (Fig. 1H). Rats preferred the chamber associated with PL-ACC activation, indicating that this pathway relieves aversion associated with peripheral hypersensitivity or allodynia (Fig. 1I-J).

Next, we examined spontaneous pain associated with inflammatory pain^12,25^. In this experiment, rats were placed in the two chambers, and only one of the chambers was paired with PL-ACC photoactivation (Fig. 1K). The conditioning stage lasted 60 minutes to allow time for spontaneous pain to occur while subjects were in the chamber. After conditioning, rats preferred the chamber associated with PL-ACC activation (Fig. 1L-M), indicating that this projection likely reduced spontaneous pain associated with persistent or chronic pain.

### Inhibition of the PL-ACC pathway reinforces pain aversion

Having shown that activation of the PL-ACC pathway is sufficient to inhibit pain aversion, we tested whether inhibiting this pathway would have the opposite effect. The evidence of such an effect would suggest the presence of a basal, tonic modulatory influence from the PL to the ACC during nociceptive processing. To do this, we injected anterograde CaMKII-NpHR bilaterally into the PL, followed by bilateral optical fiber implantations in the ACC, allowing us to completely block the synaptic connections between PL and ACC neurons (Fig. 2A). We then performed conditioned place aversion (CPA) tests. Again, rats were placed into the two-chamber device, where one of the chambers was paired with yellow light inhibition of the PL-ACC projection, whereas the other chamber was not. First, to test aversive response to acute pain in naïve rats, we used PP to trigger pain in both chambers (Fig. 2B). We found that rats strongly avoided the chamber associated with PL-ACC inhibition, indicating that blocking this pathway increased the aversive value of the noxious stimulus (Figs. 2C-E). Next, we tested the impact of inhibiting this pathway on spontaneous pain in the CFA model (Fig. 2F). After conditioning with or without optogenetic inhibition of the PL-ACC projection, CFA-treated rats avoided the chamber paired with light treatment, suggesting that inhibition of this pathway enhances the aversive value of spontaneous pain (Figs. 2G-H). These findings show that silencing the inhibitory PL inputs to the ACC enhances pain aversion and amplifies both acute and chronic pain behaviors. These data indicate a basal, tonic modulatory influence of the PL on the ACC in endogenous pain regulation and support a hierarchical organization in which the PL functions upstream to regulate pain affect.

**Figure 2:**
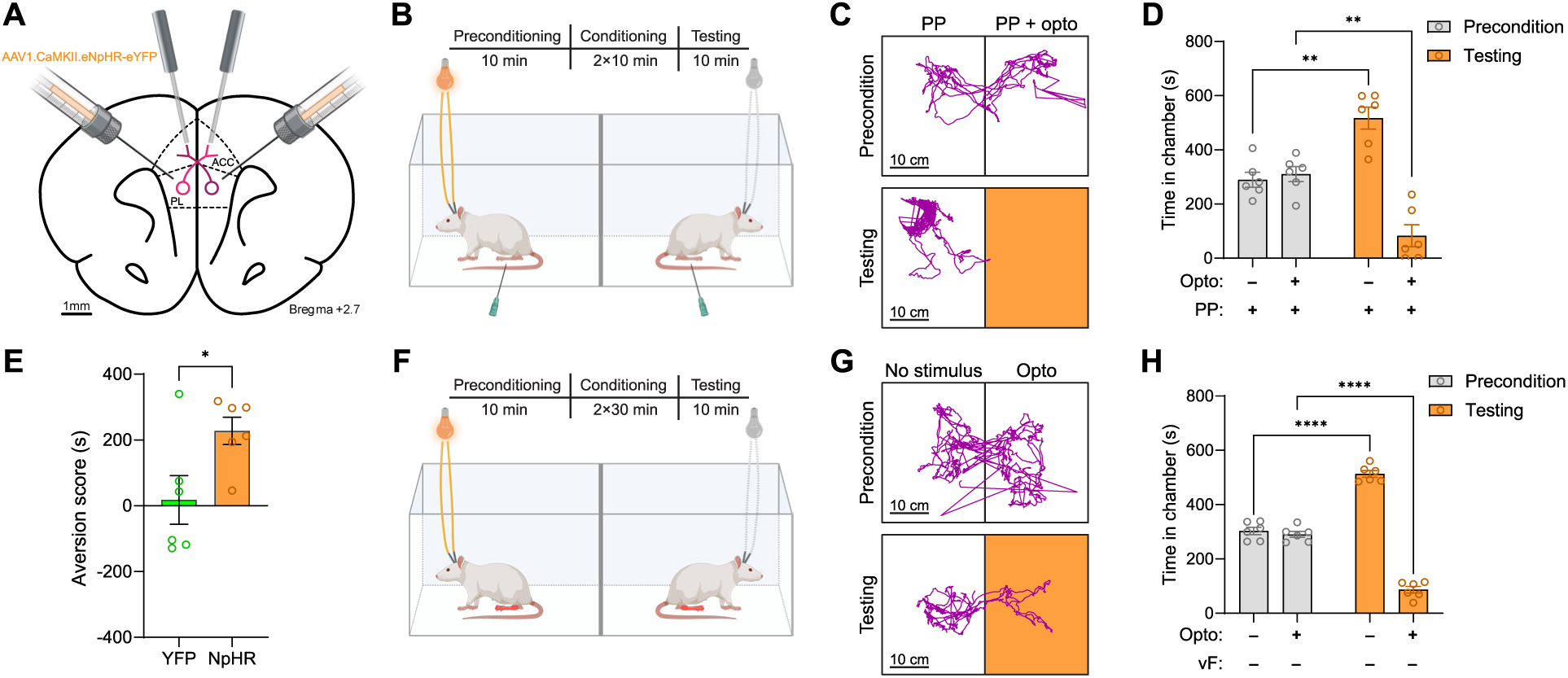
PL-ACC inhibition reinforces pain-aversive behavior. **A**Inhibitory viral injection protocol and optical fiber implantation. **B** CPA setup and experimental protocol for PP-paired conditions in acute pain model. **C-D** In the presence of acute PP, subjects demonstrate aversion toward the chamber associated with optoinhibition of PL-ACC (panel **C**: representative locomotion traces; panel **D**: p = 0.0028, n = 6, paired t-test). **E** Bilateral PL-ACC optoinhibition increases aversion scores compared to YFP controls (p = 0.0384, n = 6, unpaired t-test w/ Welch’s correction). **F** CPA setup and experimental protocol for no-stimulus conditions in chronic pain model. **G-H** In the absence of acutely noxious stimuli, chronic pain subjects demonstrate aversion toward the chamber associated with optoactivation of PL-ACC (panel **G**: representative locomotion traces; panel **H**: p < 0.0001, n = 6, paired t-test).

### PL inputs synapse onto ACC interneurons to provide feedforward inhibition of ACC outputs

To understand how PL inhibits ACC neurons, we utilized *ex vivo* patch clamp electrophysiology to identify a synaptic mechanism. We injected an anterograde AAV expressing CaMKII-ChR2-eYFP into the left PL (Fig. 3A, left panel), and subsequently prepared acute slices of right ACC that exhibited robust expression of ChR2-eYFP in PL axons (Fig. 3A, right panel). Both pyramidal neurons (Fig. 3B) and interneurons (Fig. 3C) in ACC were targeted for whole cell intracellular recording, identified post hoc based on biocytin-visualized morphology. Photoactivation of PL axons within ACC induced both excitatory and inhibitory postsynaptic potentials (EPSCs, IPSCs) in ACC pyramidal neurons (Figs. 3D, F) and interneurons (Figs. 3E, G). Analysis of response latencies from photoactivation indicated a statistically significant delay in the onset of IPSCs in both ACC pyramidal neurons (Fig. 3H) and interneurons (Fig. 3I). These results suggest that activation of PL can activate ACC interneurons and induce feedforward inhibition onto ACC pyramidal neurons.

**Figure 3:**
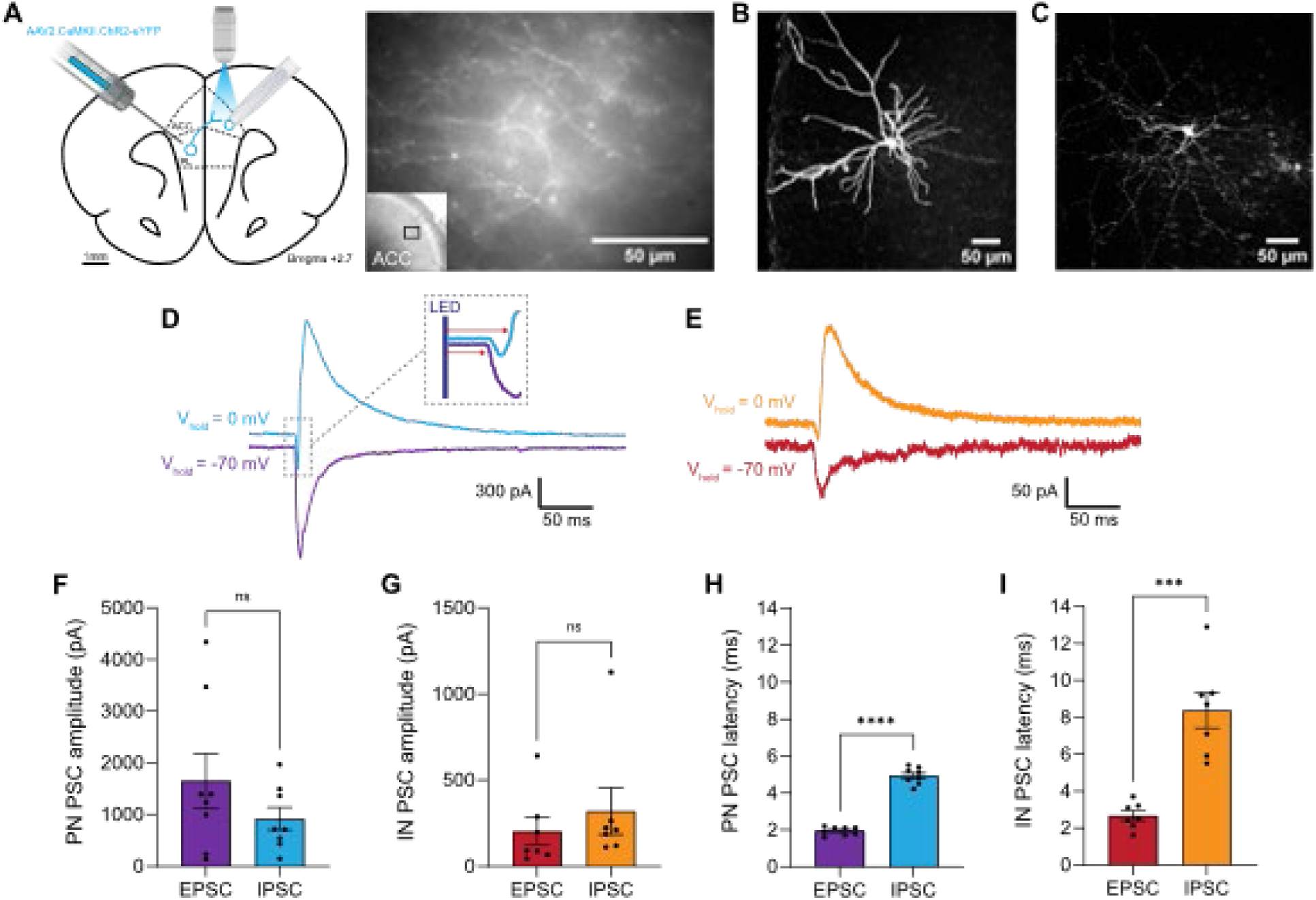
PL induces feedforward inhibition of ACC pyramidal neurons. **A** (**Left**) Excitatory viral injection protocol. (**Right**) Image of PL axons in ACC. **B** Biocytin-filled ACC pyramidal neuron. **C** Biocytin-filled ACC interneuron. **D** In an ACC pyramidal neuron, an example of excitatory postsynaptic current (EPSC, downward, V_hold_ at -70mV) and inhibitory synaptic current (IPSC, upward, V_hold_ at 0mV). Inset is an enlarged view of the outlined area, showing timing of LED-induced photoactivation and onset of EPSC and IPSC. **E** In an ACC interneuron, an example of an EPSC (downward, V_hold_ at -70mV) and IPSC (upward, V_hold_ at 0mV). **F** Amplitude of EPSCs and IPSCs in ACC PNs (n = 8). **G** Amplitude of EPSCs and IPSCs in ACC INs (n = 7). **H** Latency of EPSCs and IPSCs in ACC PNs (n = 8), p < 0.0001 by paired t-test. **I** Latency of EPSCs and IPSCs in ACC INs (n = 7), p = 0.0007 by paired t-test.

### PL activation inhibits population-level nociceptive response in the ACC

Next, to assess the role of hierarchical PL control of ACC function at the cellular level, we injected anterograde CaMKII-ChrimsonR (a red-shifted channelrhodopsin) into the left PL, followed by an injection of anterograde CaMKII-GCaMP6f within the right ACC (avoiding ipsilateral injections to minimize cross-contamination). A microendoscopic lens was then implanted into the right ACC (Fig. 4A). GCaMP6f fluorescence could thus be used as a readout of ACC pyramidal activity, while simultaneously activating axon terminals of the PL neurons in the ACC with red light. We measured Ca^2+^ activity in the ACC before and after a series of three events: optogenetic activation, noxious stimulation with a pinprick (PP) of the contralateral hindpaw, or both (Fig. 4B). Raw imaging videos were processed to identify fluorescent regions of interest (ROIs); the brightness of each ROI produced a raw fluorescence trace, which could be further processed with deconvolution to extract inferred spike times (Figs. 4C-E) using well-established processing pipelines^61,62^. To measure changes in cellular activity over time, raw fluorescence traces were used to directly calculate pre- and post-event mean fluorescence across an event-captured window of ±5 s, while inferred spike times were averaged over the same window to produce a pre- and post-event mean firing rate. First, we found that after photoactivation of the axon terminals of PL neurons, there was a significant decrease in both the mean fluorescence and firing rates of ACC neurons in the absence of noxious input (Figs. 4F-H), suggesting that PL inputs inhibit general ACC population-level activity at baseline. Next, we measured population response of excitatory neurons in the ACC to PP, with or without the activation of the presynaptic inputs from the PL (Fig. 4I). As expected, ACC neurons increased their firing rates in response to the noxious stimulus^12,23^. However, optogenetic activation of the presynaptic PL inputs partially reversed the increase in ACC activity ordinarily associated with acute pain (Figs. 4J-K).

**Figure 4:**
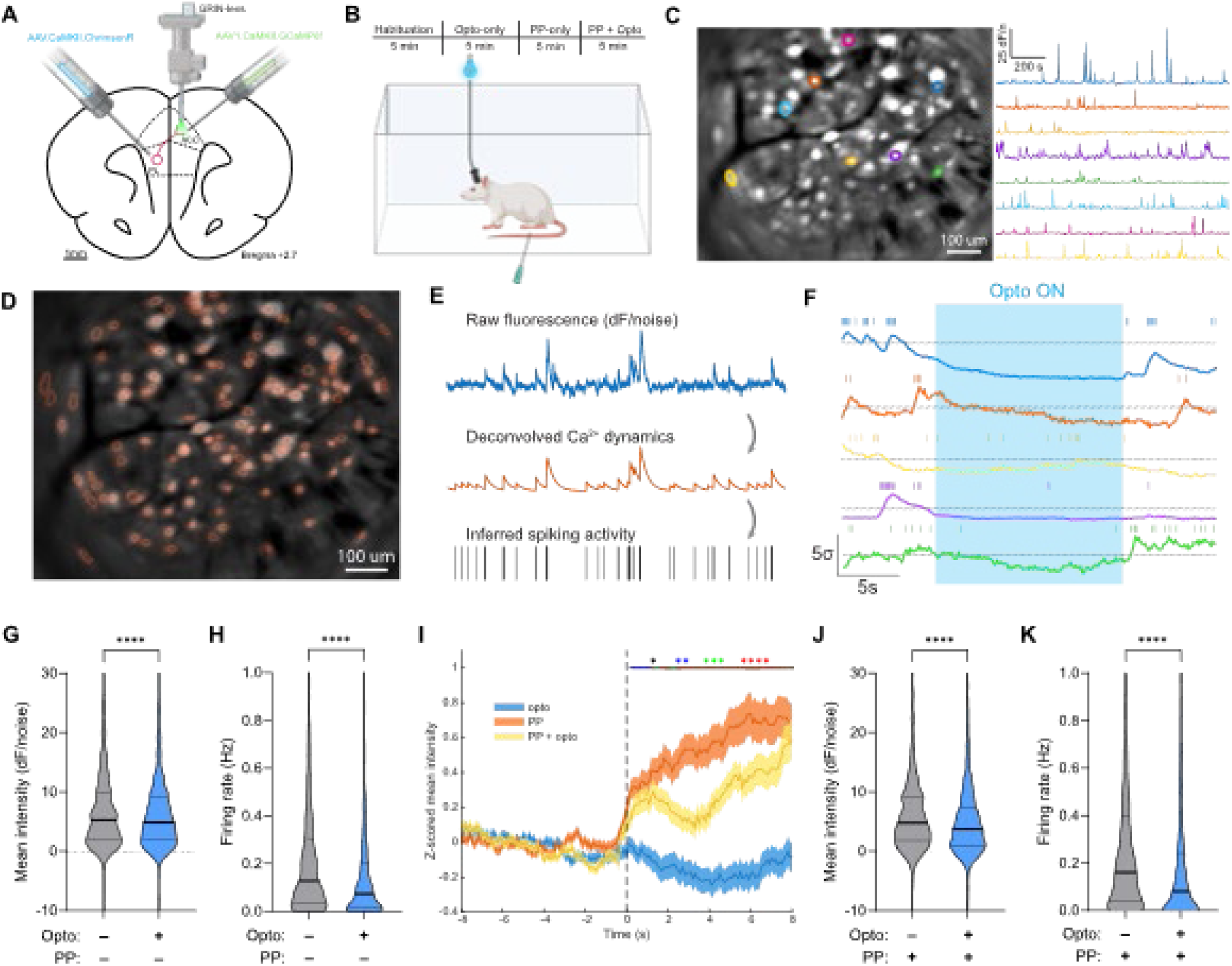
PL inputs inhibit the ACC response to noxious stimuli. **A** Viral injection protocol and GRIN-lens implantation. AAV.CaMKII.ChrimsonR was injected into the left PL, and a GRIN lens was implanted into the right ACC following injection of AAV1.CaMKII.GCaMP6f. **B** Ca^2+^ imaging setup and experimental protocol. **C** Eight representative cellular fluorescence traces, extracted from imaging field of view. **D** Map of all detected cellular contours overlaid on maximum-intensity projection image. **E** Overview of deconvolution and spike inference postprocessing. **F** Five representative peri-opto raw fluorescence traces, normalized to the pre-opto period, with inferred spikes overlaid as ticks. **G-H** Neurons show decreased post-event mean fluorescent intensity (**G**, n = 872, p < 0.0001, Wilcoxon signed-rank test) and firing rate (**H**, n = 872, p < 0.0001, Wilcoxon signed-rank test) following PL optoactivation. **I** Peri-event mean Ca^2+^ fluorescence Z-scores for opto, PP, and PP+opto conditions, among all neurons. (n = 872, * < 0.05, ** < 0.01, *** < 0.001, **** < 0.0001, per-timepoint one-way ANOVA). **J-K** ACC neurons show decreased post-event mean fluorescent intensity (panel **J**, n = 872, p < 0.0001, Wilcoxon signed-rank test) and firing rates (panel **R**, n = 872, p < 0.0001, Wilcoxon signed-rank test) when PP is paired with PL optoactivation.

### PL activation gates nociceptive information transmission in the ACC

Next, we examined the network-level effects of PL gating on ACC function. The nociceptive network connectivity within the ACC is driven by specific pain-responsive neurons – neurons that significantly increased their activity in response to painful stimuli^12,23^. Thus, we first examined how PL inputs inhibit these pain-responsive ACC neurons. Prior studies have shown that approximately 15-20% of ACC neurons are pain-responsive^12,23^; we confirmed this finding (Fig. 5A). Optogenetic activation of the axon terminals of PL neurons in the ACC decreased the baseline activity of this select population of neurons in the absence of noxious stimuli (Figs. 5B-C). We Z-scored each smoothed fluorescence trace relative to the pre-PP baseline, enabling comparison of post-event mean and peak fluorescence, as well as firing rate, across treatment conditions. We found that optogenetic activation of the presynaptic inputs from PL significantly reduced nociceptive-evoked population activity among ACC pain-responsive neurons (Figs. 5D-E). We then quantified neuronal responsiveness using two complementary activity metrics to capture multi-dimensional modulation. First, the mean response intensity reflects sustained neuronal engagement, providing insight into prolonged coding rather than transient peak bursts. Its reduction suggests that modulatory input from the PL persists throughout nociceptive processing in the ACC (Fig. 5F). In addition, a decrease in inferred firing rates of ACC neurons reinforces that the underlying spike dependent activity is suppressed, not merely calcium signal properties (Fig. 5G). Together, these results indicate that PL inputs have a strong inhibitory effect on the ability of ACC excitatory neurons to respond to noxious stimuli.

**Figure 5:**
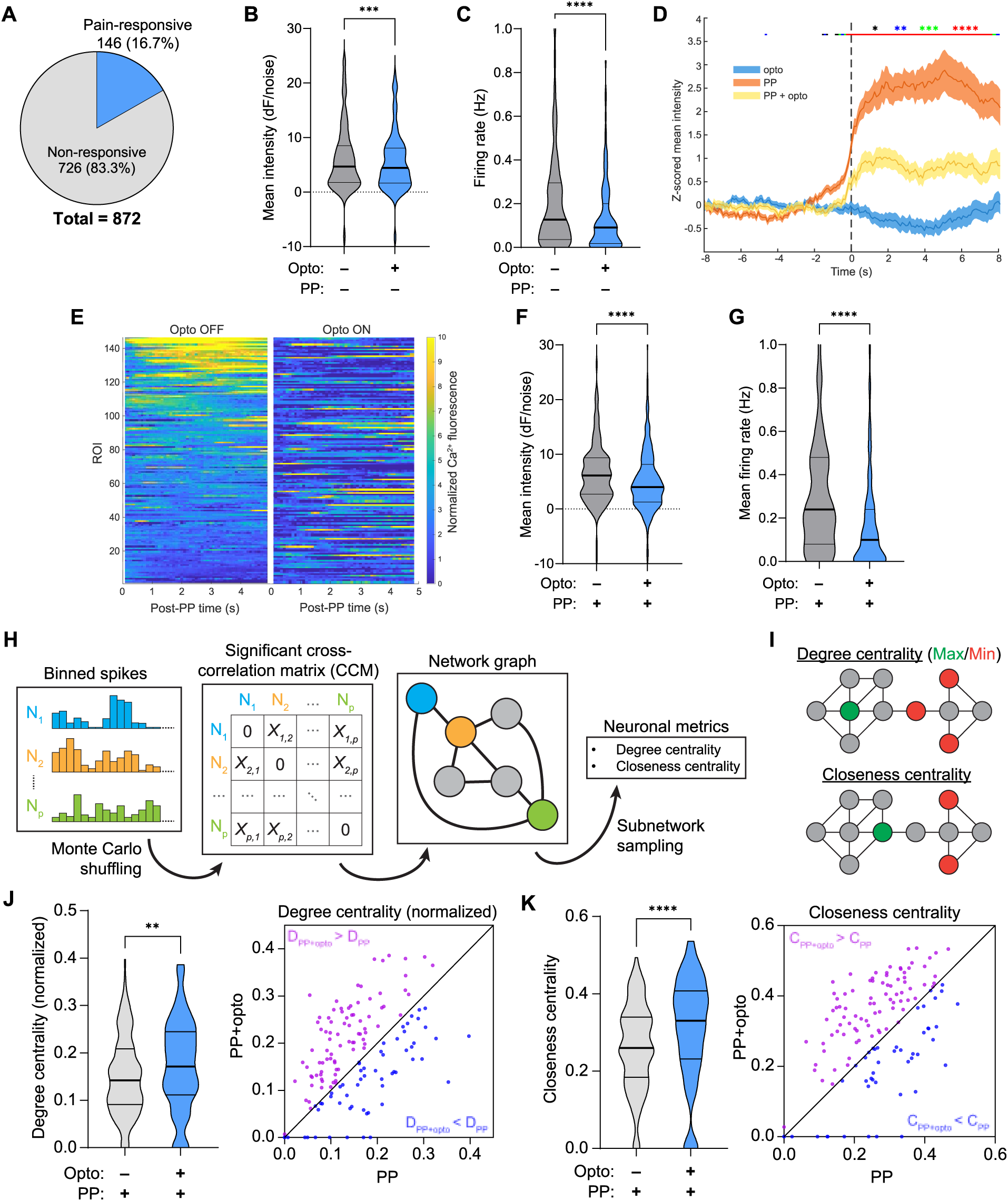
PL induces functional centralization of pain-responsive ACC neurons. **A** Proportion of ACC neurons responsive to PP (“pain-responsive”). **B-C** Pain-responsive neurons show decreased post-event mean fluorescent intensity (panel **B**, n = 146, p = 0.0001, Wilcoxon signed-rank test) and firing rates (panel **C**, n = 146, p = 0.0001, Wilcoxon signed-rank test) from baseline following PL optoactivation. **D** Peri-event mean Ca2+ fluorescence Z-scores for opto, PP, and PP+opto conditions, among pain-responsive neurons. (n = 146, * < 0.05, ** < 0.01, *** < 0.001, **** < 0.0001, per-timepoint one-way ANOVA). **E** Post-PP Z-scored Ca2+ fluorescence, averaged across all trials for pain-responsive neurons (n = 146), in the absence (left) and presence (right) of PL optoactivation; neurons are aligned from high to low fluorescence in the absence of optoactivation, and consistent across rows. **F-G** Pain-responsive neurons show decreased post-event mean fluorescent intensity (panel **F**, n = 146, p < 0.0001, Wilcoxon signed-rank test), and firing rates (panel **G**, n = 146, p < 0.0001, Wilcoxon signed-rank test) when PP is paired with PL optoactivation. **H** Overview of graph-theoretic analysis. **I** Visualization of degree and closeness centrality metrics in an example graph. **J** Pain-responsive neurons show increased degree centrality (n = 146, p = 0.0011, Wilcoxon signed-rank test) in the acute pain setting when PP is paired with PL optoactivation. Violin plots (**left**) show aggregate data, while scatter plot (**right**) shows individual unit degree with respect to the identity line. **K** Pain-responsive neurons show increased closeness centrality (n = 146, p < 0.0001, Wilcoxon signed-rank test) in the acute pain setting when PP is paired with PL optoactivation. Violin plots (**left**) show aggregate data, while scatter plot (**right**) shows individual unit degree with respect to the identity line.

We next investigated how inhibitory PL inputs modulate overall network activity in the ACC through their modulation of these pain-responsive neurons. To do this, we used graph-theoretic analysis to characterize functional connectivity and information transmission among ACC neurons^28,44^. We first inferred putative spike trains from preprocessed prefrontal Ca²⁺ fluorescence signals using a nonnegative deconvolution approach (Fig. 4E). These inferred nonnegative activity traces were then used to calculate scale-invariant cross-correlations between neuronal ROIs^45^, enabling construction of functional graphs in which nodes corresponded to individual neurons and edges reflected significant correlation strengths (Fig. 5H).

We applied an established method to compute two graph-network statistics: degree centrality and closeness centrality, to test the functional coordination of pain-responsive neurons in processing nociceptive information within the ACC^28,44^. Degree centrality quantifies the number of neighboring nodes, reflecting changes in functional connectivity, while closeness centrality measures how close a node is relative to all other nodes, thereby indicating its influence in information integration across the network (Fig. 5I). We found that after PL activation, there was a significant increase in both degree centrality (Fig. 5J) and closeness centrality (Fig. 5K) in pain-responsive neurons. These findings indicate that PL activation increases the centralization of nociceptive information flow within the ACC. This signal centralization, in the context of reduced excitability at the single-neuron level, suggests that the ACC network enters a gated, low-output state, where only a diminished subset of neurons drive the nociceptive response in the ACC, and they do so less frequently (Fig. 6).

**Figure 6:**
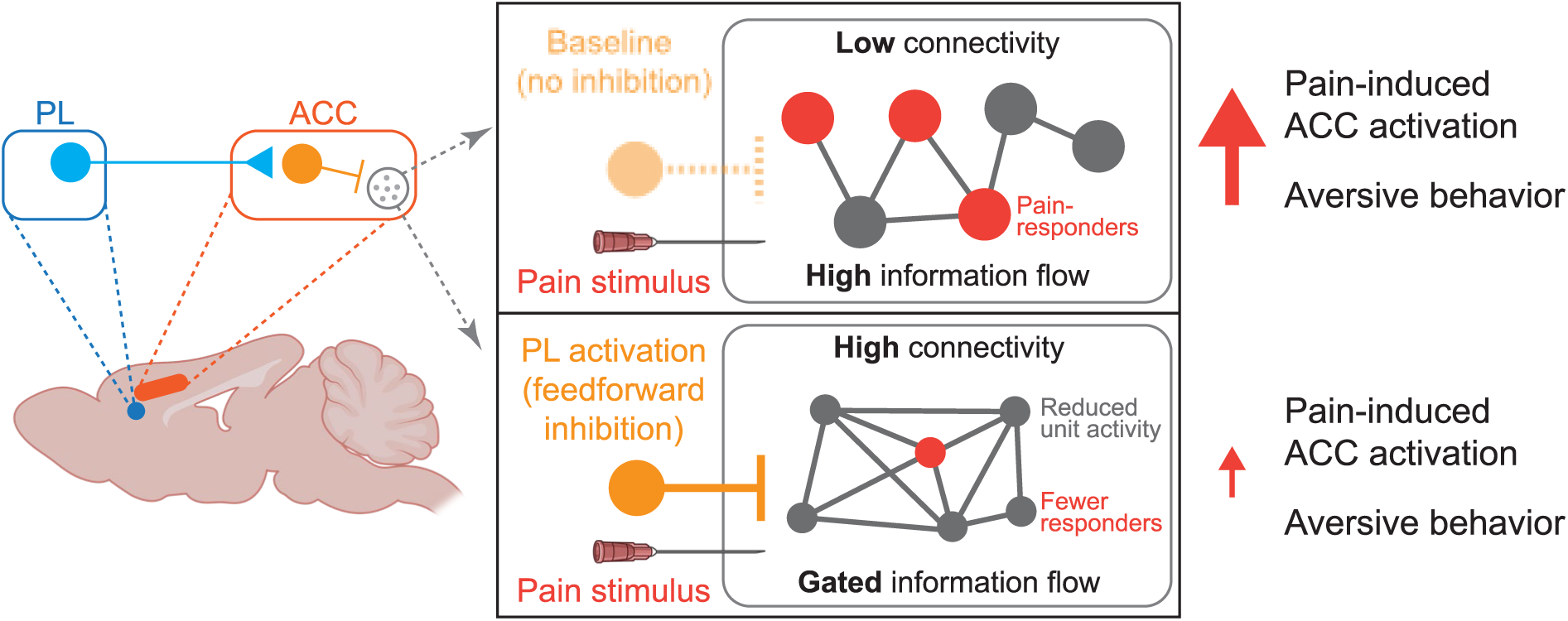
Graphic summary of PL-ACC effects on ACC functional connectivity and pain response. PL inputs modulate pain-induced ACC activation and aversive behavior through three related mechanisms. Most directly, PL projections synapse to ACC interneurons to produce feedforward inhibition of ACC activity. This inhibition induces a centralization of ACC pain-responsive neurons, which serves to gate information flow. This information gating reduces global ACC neuronal activity and downstream pain-aversive behavior.

## Discussion

The PFC and the ACC are central nodes in the cortical processing of the sensory and affective dimensions of pain. Although they are reciprocally connected anatomically, a functional circuit linking these regions in pain regulation has remained undefined. In this study, we demonstrate that PL neurons project directly to the ACC to inhibit its excitatory outputs ^62^. This PL-mediated inhibition of ACC activity reduces pain-aversive behaviors in both acute and chronic pain states. These findings establish a PL–ACC circuit that acts as a hierarchical gate controlling pain-related behaviors.

Our study provides a direct circuit mechanism by which the PFC, a critical region for endogenous sensory regulation, modulates ACC activity. This mechanism is supported by several independent lines of evidence. First, histology studies confirm the existence of this direct anatomic pathway^43,50^. Second, optogenetic activation of PL axon terminals decreased the output of ACC excitatory neurons. Third, graph-theoretical analysis indicates that PL activation results in a bottleneck of nociceptive information gating in the ACC. Fourth, *ex vivo* patch clamp recordings indicate that activation of the PL axons in the ACC can excite ACC interneurons and induce feedforward inhibition of ACC pyramidal neurons. Lastly, behavioral experiments demonstrate that activation of this projection reduces pain aversion, whereas its inhibition enhances aversion, in animal models of both acute and chronic spontaneous pain.

Similar forms of feedforward cortico-cortical inhibition have been proposed to regulate processes such as fear conditioning and preparatory attention^51–53^. Our findings here therefore raise the possibility that, in addition to its well-characterized direct cortico-subcortical projections^33,36–38,54^, the PFC may also modulate pain and other sensory–affective experiences via gating of the ACC. Moreover, our behavioral experiments, particularly those involving photoinhibition, strongly indicate a baseline, tonic inhibitory influence from the PL to the ACC. This tonic inhibition suggests a cortical hierarchy in the regulation of affective experience, in which the PFC exerts modulatory influence on ACC nociceptive processing. At the mechanistic level, this tonic inhibition is characterized by reduced firing rates in pain-responsive neurons that nonetheless occupy more central positions within the functional network, yielding a gated or bottlenecked state with low overall output. At a synaptic level, such inhibition could be mediated by the PL engagement of local parvalbumin- and/or somatostatin-expressing interneurons, which have been shown to produce feedforward inhibition of ACC pyramidal neurons^55^.

Chronic pain is associated with increased plasticity in the ACC but hypoactivity in the PFC^12,13,29–32,39–41,46,56,57^. Previous studies have focused on their parallel subcortical projections to explain how this imbalance in the activity levels of PFC and ACC contributes to chronic pain. Our study supports a direct circuit mechanism underlying this imbalance: reduced pyramidal outputs from the PFC may diminish tonic inhibitory inputs to the ACC, leading to increased excitatory output from ACC neurons.

Several recent studies have revealed functional connections between the ACC and the primary somatosensory cortex (S1) and insular cortex^23,58–60^. In these studies, the S1 and insular inputs increase ACC activity, leading to enhanced pain aversion. Together with our results, these studies indicate that the ACC may function as a central hub for aversive processing, regulated by a complex network of cortical projections to shape behavioral responses to pain^61^.

Future studies could build on these findings by further elucidating ACC microcircuit dynamics in response to PL gating. Because our work in rats limits the resolution with which specific interneuronal populations can be examined, investigations in mice may offer deeper insight into the complex interactions between the PL and ACC. Extending this line of research to humans could also clarify the analogous functional connectivity between the ACC and the human homolog of the PL, the dorsolateral prefrontal cortex.

In summary, we identify a novel pain-regulatory cortico-cortical pathway from the PFC to the ACC in which the PFC acts as a top-down gate within a hierarchical cortical network. This hierarchical circuit provides a key mechanistic link in cortical pain regulation and represents a promising target for neuromodulatory interventions in chronic pain and related neuropsychiatric disorders.

## Methods

### Animals

All procedures were performed in accordance with New York University Grossman School of Medicine (NYUGSOM) Institutional Animal Care and Use Committee (IACUC) guidelines to minimize animal use and discomfort, as consistent with the NIH *Guide for the Care and Use of Laboratory Animals*. Genetically wild-type male Sprague-Dawley rats were purchased from Taconic Biosciences (NTac:SD), and pair-housed at the vivarium in the NYU Langone Science Building. Rats were housed in humidity and temperature-controlled conditions, under a 12-hour (6:30 AM-6:30 PM) light-dark cycle. Vendor health reports confirmed that the rats were free of known pathogens. All rats were received at a developmental stage of 8 weeks and given approximately 10 days to adjust to their new environment prior to any experimentation. All rats were naive to procedures and drugs before surgical procedures.

### Drugs

0.1 mL of Freund’s complete adjuvant (CFA) (inactivated *Mycobacterium tuberculosis*, Sigma-Aldrich) was suspended in a 1:1 emulsion with saline and injected subcutaneously into the plantar aspect of the left hindpaw to induce inflammatory pain.

### Viral construction and packaging

Recombinant adeno-associated viral (AAV) vectors were serotyped with AAV coat proteins and packaged at Addgene viral vector manufacturing facilities. Viral titers contained approximately 5 × 10^12^ particles per milliliter for pENN.AAV1.CamkII.GCaMP6fWPRE.SV40, pAAV1.-CaMKIIa.-eNpHR.3.0.EYFP, AAV2.CamKIIa.hChR2(H134R).eYFP.WPRE.hGH, pAAV.- CamKIIa.ChrimsonR.mScarlet.KV2.1, and AAV1.CaMKII(1.3).eYFP.WPRE.hGH. Aliquots were stored in a light-protected freezer before use.

### Intracranial viral injections

Consistent with previous experiments^36^, rats were anesthetized with isoflurane (1.5-2%). Viruses were delivered to the PL-PFC in all animals, and additionally in the ACC for calcium imaging subjects. For the PL-PFC, rats were unilaterally or bilaterally injected with 0.7 µL viral vectors at a rate of 0.1 µL/20 s with a 32-gauge 1 µL Hamilton syringe at anteroposterior (AP) +2.9 mm, mediolateral (ML) ±1.6 mm, and dorsoventral (DV) -3.5 mm, with the tips angled 17° toward the midline. For the ACC, rats were unilaterally or bilaterally injected with 0.7 mL viral vectors at a rate of 0.1 µL/20 s with a 32 G 1 µL Hamilton syringe at AP +2.9 mm, ML ±1.6 mm, and DV - 2.0 mm, with the tips angled 22° toward the midline. After injecting the full volume, microinjection needles were left in place for 10 min before being raised 0.5 mm, thereby allowing for diffusion of viral particles and minimization of reflux along the injection tract. The microneedle was left in place for an additional 2 min before being slowly raised out of the brain. Immediately following completion of injections, implantations of GRIN-lenses or optical fibers were conducted as described in the following sections. Rats were allowed to recover for 4-6 weeks post-injection to provide time for adequate viral expression prior to initiation of experiments.

### Gradient-index lens implantation

As in previous studies^28,44^, immediately following viral injection, rats used for calcium imaging were stereotactically implanted with an integrated gradient-index (GRIN) lens and baseplate (1.0 mm diameter, 9.0 mm length, Inscopix) at AP +2.9 mm, ML ±1.6 mm, and DV -1.8 mm, with the tips angled at 22° toward the midline, placing the lens +100-300 µm above the imaging plane. The gap between the placement of the lens and the opening of the craniotomy site was filled with silicone elastomer (Kwik-Sil, World Precision Instruments). Integrated lenses were permanently affixed to the skull with acrylic dental cement.

### Optical fiber implantation

Prior to implantation, 200 µm optical fibers were secured in 2.5 mm ceramic ferrules (ThorLabs). Immediately following viral injection, rats used for behavioral testing were stereotactically implanted with unilateral or bilateral fibers in the ACC (respective to the PL viral injections) at AP +2.9 mm, ML ±1.6 mm, and DV -1.5 mm, with the tips angled at 22° toward the midline. Fibers with ferrules were permanently affixed to the skull with acrylic dental cement.

### Calcium imaging procedure

For each imaging session, the subject was placed within a cage-sized recording chamber (∼30 × 45 cm) over a mesh table, as reported previously^12^, and a miniature microscope (nVoke, Inscopix) was secured to the integrated baseplate. A wired webcam linked to the recording computer was used to record the experiments, and a pulsed LED was used to synchronize events across both recordings. Recordings began with a baseline/habituation period, during which the animal was allowed to freely move within the testing box, without any external stimuli. Following habitation, all recording sessions consisted of three stages: opto-only, PP-only, and PP + opto. Each stage contained approximately 5 trials, with variable inter-trial intervals of ∼60 s to avoid sensitization, and inter-stage intervals of ∼180 seconds to allow Ca^2+^ signals to return to baseline. Red light was delivered at 20mW to excite ChrimsonR through the GRIN-lens during opto stages, at a frequency of 20 Hz with 20% duty cycle and 15s stimulation durations. For pinprick (PP) stages, noxious pinprick stimulation was applied by lightly pricking the plantar surface of the hind paw contralateral to the brain recording site with a 27-gauge needle through the mesh table. Noxious stimulation was terminated by withdrawal of the paw, which occurred in all cases. No behavioral sensitization or physical damage to the paws was observed. For PP + opto conditions, PP was timed for delivery at least 5 s into each stimulation window.

### Calcium fluorescence data acquisition

As reported previously^28^, all fluorescent microscope videos were recorded with a fluorescence power of 0.5-0.8 mW/mm^2^ at 1280 x 800 pixels and a frame rate of 10 Hz. Following acquisition, raw videos were spatially downsampled by a binning factor of 4 (16x spatial downsampling) using Inscopix Data Processing Software (IDPS, Inscopix). Next, videos were motion-corrected with reference to a single frame using IDPS to minimize motion artifacts. Ca^2+^ signals were then extracted using the constrained non-negative matrix factorization (CNMFe) algorithm^62,63^, as implemented in IDPS. This process estimates spatiotemporally constrained instances of calcium activity for each neuron relative to background noise to produce a single dF/noise trace for each region of interest (ROI). Finally, IDPS was used to infer spike times and produce neuronal ROI contour maps, which were exported as .CSV and .TIFF files respectively for further analysis.

### Calcium response analyses

Calcium fluorescence traces were processed for each neuron using a 500 ms moving average window, which was used to reduce the effect of signal transients from motion artifacts. As outlined previously^28^, trial-averaged Ca^2+^ fluorescence traces were then obtained for individual neurons by capturing an event-triggered window of -5 to 5 s, where 0 denotes the event time (i.e., vF, PP, or opto-ON time for opto-only). Each point in a trial was Z-scored using the values within the baseline period of -4.5 to -1.5 s prior to each event. Z-scored trials for a single neuron were then averaged to produce a single mean Ca^2+^ fluorescence trace per condition per neuron. Pre-event mean fluorescence was calculated as the overall mean from -5 to 0 s, while post-event mean was calculated from 0 to 5 s.

Mean firing rate was obtained by capturing all inferred spike times within an event-triggered window of -5 to 5s. Total pre-event spikes (within the -5 to 0 s window) and post-event spikes (within the 0 to 5 s window) were counted and then divided by 5 seconds to produce a single pre-and post-event mean firing rate per trial. These were further averaged across trials to produce a pre- and post-event mean firing rate per condition per neuron. Finally, each set of values was aggregated for each neuron among all rats for inter-condition analysis.

### Graph-theoretical analysis

An adaptation of previously reported methods^28,44^. For all recorded units in a calcium imaging session, inferred spiketimes from a 300 s recording window (beginning at the start of the recording for the baseline condition, and immediately after the first event sample for opto, PP, and PP+opto conditions) were counted into 200-ms bins. Next, for each pair of units, the maximum cross correlation within a lag period of ±1 s was calculated. To identify significance thresholds for these cross-correlations, we applied a Monte Carlo simulation method, where one unit of each pair was randomly shuffled to form a distribution of maximum correlogram values across 1000 iterations. True cross-correlation values were deemed significant if they fell above the 95^th^ percentile within this distribution; these significant values were then arranged into an adjacency matrix, which was used to form an undirected network graph for each recording session.

Comparison of graph-theoretic metrics between pain-responsive units across sessions requires correcting for the differing sizes of each network, as well as the proportions of pain-responsive units. To achieve this, for each session, we sampled nodes randomly to produce bootstrapped subgraphs matching the size of the session with the smallest network, with a proportion of pain-responding nodes matching the minimum pain response rate across all sessions (rounded to the nearest integer). Degree and closeness centrality were calculated for each subgraph, with normalized degree for a subgraph of size N calculated by dividing by (N-1). This was performed up to a maximum of 1000 iterations for each session; however, if the number of unique subgraphs in a session was less than 1000, only one iteration per unique subgraph was included, to avoid oversampling. Graph metrics were then averaged across all iterations for each unit to produce the final datasets.

### Immunohistochemistry

Individual ACC neurons were filled via patch pipette with 0.2% biocytin during whole cell recordings and visualized by adapting a published biocytin immunohistochemistry protocol^64^. After recording, slices were fixed overnight in 4% paraformaldehyde (PFA) in 1x PBS. Slices were washed in 1x PBS (3 × 10 min), permeabilized, and blocked with 10% normal goat serum and 0.3% Triton X-100 in 1x PBS for 1 h. Slices were then incubated overnight at 4°C in 3% normal goat serum, 0.3% Triton X-100, and 1:1,000 streptavidin-bound Alexa Fluor 647 conjugate (S32357; Invitrogen). Slices were washed in 1x PBS (3 × 15 min), mounted in Fluoromount-G with DAPI medium (00-4959-52, Invitrogen) and imaged. Images of individual biocytin-filled neurons were captured on an upright laser scanning confocal microscope (LSM 800; Zeiss). High resolution z-stacks were taken at 20x (Plan-Apochromat 0.8 NA; 1,024 × 1,024 minimum frame resolution; 8-bit pixel depth). If spine visualization or dendrite morphology were unclear at 20x, additional images were taken at 63x (Plan-Apochromat 1.40 NA Oil DIC; 1,024 × 1,024 minimum frame resolution; 8-bit pixel depth) for confirmation. ACC excitatory pyramidal neurons and inhibitory interneurons were visually identified based on morphological architecture and complexity, somata size and dendritic spine density. Specifically, pyramidal neurons featured a larger soma, vertically projecting apical dendrites, with multiple oblique dendrites prior to distal tuft. Interneurons featured a small soma, less prominent apical dendrite, thinner dendrites, and fewer spines.

### *Ex vivo* patch-clamp electrophysiology

Acute ACC brain slices were prepared as follows. Rats were injected in the left PL-PFC with an AAV expressing hChR2 as above. After sufficient time for AAV infection, rats were deeply anesthetized with isoflurane for 12 min. After intracardiac perfusion with artificial cerebrospinal fluid (ACSF), the brain was quickly dissected in oxygenated (95% O2 and 5% CO2) dissection ACSF containing (in mM): 10 NaCl, 2.5 KCl, 0.5 CaCl2, 7 MgCl2, 1.25 NaH2PO4, 25 NaHCO3, 195 Sucrose, 10 Glucose, 2 Na-Pyruvate (pH: 7.4). Bilateral coronal brain slices (400 μm) containing the ACC were prepared with a vibratome (Leica VT-1200s, Deer Park, IL), placed in warm (34 °C) 50% dissection ACSF containing and 50% ACSF containing (in mM): 125 NaCl, 25 NaHCO3, 2.5 KCl, 1.25 NaH2PO4, 2 CaCl2, 1 MgCl2, 22.5 Glucose, 3 Na-Pyruvate, 1 Ascorbic Acid (pH: 7.4) for 15 min, and then at room temperature for at least 1 h.

Whole-cell voltage-clamp recordings were performed as follows. Right ACC slices were placed in a recording chamber with oxygenated ACSF at 32 °C. Recording pipettes (3-5 mΩ) were filled with intracellular solution (in mM): 135 Cs-Methanesulfonate, 5 KCl, 0.1 EGTA-Na, 10 HEPES, 2 NaCl, 5 ATP, 0.4 GTP, 10 phosphocreatine (pH 7.35), as well as 0.2% biocytin was included in the intracellular solution. ACC neurons were targeted for whole cell intracellular recording in voltage clamp mode to record postsynaptic currents. Pyramidal or interneuron identity were confirmed post-hoc by review of morphology via biocytin staining. During intracellular recording, PL-PFC axons were excited using 100% max intensity of a single wavelength pulse (470 nm wavelength, 2 ms pulse duration) with CoolLED (PE-4000, Andover, UK) delivered via a 60x objective placed above the slices. Excitatory and inhibitory postsynaptic currents were recorded at a holding potential of -70 mV and 0 mV, respectively. Synaptic current latencies were measured from stimulus artifact to onset of response.

### Optogenetic setup

Prior to each experiment, optic fibers were connected to a diode-pumped solid-state (DPSS) laser (Ultralasers, Inc.) through a mating sleeve, as described previously^36^. Laser output power at the fiber tip was calibrated to 6 mW. For ChR2 activation and YFP controls, blue light (460 nm) was delivered through the optic fiber using pulse protocols generated with a transistor-transistor logic (TTL) pulse generator (Doric Lenses), at a pulse frequency of 20 Hz with 20% duty cycle, and stimulations were delivered for 5 s every 15 s over the full conditioning period. For NpHR activation, yellow (589 nm) light was delivered as a continuous wave for the full conditioning period.

### Conditioned place aversion assay

Conditioned place aversion (CPA) experiments were conducted similarly to those described previously^12^. A standard two-compartment apparatus was used, consisting of two equal cage-sized chambers (∼30 × 45 cm). Different randomly selected flavors of scented petrolatum were applied to the walls of each chamber to provide contextual gusto-olfactory cues. The two compartments were separated by an opening large enough for a rat to comfortably and freely travel through and placed on top of a metal mesh table. Each CPA protocol included one preconditioning (baseline), two conditioning, and one testing phase. During preconditioning (10 min), animals were allowed to travel freely between chambers in the absence of external stimuli; animals spending > 80% of the total time in either chamber during the preconditioning phase were eliminated from further analysis. Immediately following the preconditioning phase, rats underwent conditioning (2 × 10 or 2 × 30 min), during which both chambers were paired with a common stimulus (PP, 0.6g vF, or no stimulus), while only one chamber was paired with optogenetic stimulation. The common stimulus was repeated every 15 s, and for pulsed protocols, this stimulus was timed to occur roughly halfway through the optogenetic stimulus duration. Chamber conditions and the order of chambers were counterbalanced across all rats. Finally, during the testing phase (10 min), rats were again allowed free access to both chambers without any external stimulation. Movements of the rats in each chamber were recorded by a camera and analyzed with motion-tracking software (ANY-maze). Decreased time spent in a chamber during the test phase as compared with the baseline indicated avoidance (aversion) of that chamber, whereas increased time spent in a chamber during the test phase as compared with the baseline indicated preference for that chamber. This effect was captured by an aversion score for each condition, defined as (T_preconditioning_ – T_testing_), where T indicates the time spent in the chamber associated with that condition for the subscripted experimental stage.

### Mechanical allodynia test

Mechanical allodynia was measured using a Dixon up-down method with von Frey monofilaments. Rats were placed individually into closed acrylic chambers over a mesh table and allowed to habituate for 20 mins prior to testing. vF filaments with logarithmically incremental stiffnesses were then applied to the hind paw, as described previously^65^. 50% withdrawal thresholds were calculated.

### Quantification and statistical analysis

All statistics were conducted similarly to those described previously^28^. Bar graphs with error bars are given as mean ± SEM unless otherwise stated. To compare Ca^2+^ metrics across baseline vs opto or PP vs PP+opto conditions, a two-tailed paired Wilcoxon rank-sum test was used. For the CPA assay, a two-tailed paired Student’s t-test was used to compare the time spent in each treatment chamber before and after conditioning. A two-tailed unpaired Student’s t-test with Welch’s correction was used to compare differences in aversion scores across conditions. Sample sizes were defined as to be comparable to previous studies. Exact values of n and p are reported in the figure legends.

To identify neurons that altered their firing rate in response to acute PP (termed “PP-responsive”), we adapted a previously described method^28^. For each trial, event-triggered windows of raw fluorescence from -5 to 5s were gathered and z-scored using values in the baseline range of -4.5 to -1.5 s. The mean fluorescence of the 2 s window from -5 to -3 s was used as the pre-stimulus value. The mean fluorescence of the 5 s window from 0 to 5 s was used as the post-stimulus value. A one-tailed Wilcoxon rank-sum test was then performed between pre- and post-stimulus values over all PP-only trials, and the neuron was considered PP-responsive if the mean increased post-PP with p < 0.05.

Type I error rate (α) was set at 0.05 for all hypothesis testing unless otherwise stated. GraphPad Prism 10 (GraphPad Software) and MATLAB R2025b (MathWorks) were used for all statistics and figure generation.

## Acknowledgements

This study was primarily funded by the National Institute of General Medical Sciences (NIGMS) federal research grant R01-GM115384, with additional support from the National Institute of Neurological Disease and Stroke grant R01-NS123928 and the Interdisciplinary Pain Research Program at NYU Grossman School of Medicine. Z.S.C. was additionally supported by federal research grants RF1-DA056394, P50-MH132642, and R01-MH139352 from the US National Institute of Health.

## Author contributions

J.W., Q.Z. and E.H., designed the experiments; E.H., G.S., E.Z., and Q.Z. performed all surgical preparations; E.H., G.S., and E.Z. performed calcium imaging, behavioral experiments, and statistical analyses; E.H. conducted graph-theoretical analysis under the direction of Q.Z and Z.S.C.; C.T., I.R., and A.V.M. performed patch-clamp electrophysiology and histological analysis. J.W., Q.Z., E.H., and A.V.M. wrote the manuscript.

## Ethics declarations

### Data Availability

The authors declare that all the data supporting the findings of this study are available within the paper.

### Code Availability

Custom MATLAB code are available upon request.

### Materials and correspondence

Please direct all correspondence and material requests to.

## References

1 Salzman, C. D. & Fusi, S. Emotion, cognition, and mental state representation in amygdala and prefrontal cortex. Annual review of neuroscience 33, 173–202 (2010). 10.1146/annurev.neuro.051508.135256

2 Miller, E. K. The prefrontal cortex: complex neural properties for complex behavior. Neuron 22, 15–17 (1999). 10.1016/s0896-6273(00)80673-x

3 Apkarian, A. V., Bushnell, M. C., Treede, R. D. & Zubieta, J. K. Human brain mechanisms of pain perception and regulation in health and disease. European journal of pain 9, 463–484 (2005). 10.1016/j.ejpain.2004.11.001

4 Talbot, J. D. et al. Multiple representations of pain in human cerebral cortex. Science 251, 1355–1358 (1991).

5 Bushnell, M. C., Ceko, M. & Low, L. A. Cognitive and emotional control of pain and its disruption in chronic pain. Nature reviews. Neuroscience 14, 502–511 (2013). 10.1038/nrn3516

6 Bliss, T. V., Collingridge, G. L., Kaang, B. K. & Zhuo, M. Synaptic plasticity in the anterior cingulate cortex in acute and chronic pain. Nature reviews. Neuroscience 17, 485–496 (2016). 10.1038/nrn.2016.68

7 Rainville, P., Duncan, G. H., Price, D. D., Carrier, B. & Bushnell, M. C. Pain affect encoded in human anterior cingulate but not somatosensory cortex. Science 277, 968–971 (1997).

8 Johansen, J. P., Fields, H. L. & Manning, B. H. The affective component of pain in rodents: direct evidence for a contribution of the anterior cingulate cortex. Proceedings of the National Academy of Sciences of the United States of America 98, 8077–8082 (2001). 10.1073/pnas.141218998

9 LaGraize, S. C., Borzan, J., Peng, Y. B. & Fuchs, P. N. Selective regulation of pain affect following activation of the opioid anterior cingulate cortex system. Experimental neurology 197, 22–30 (2006). 10.1016/j.expneurol.2005.05.008

10 Navratilova, E. et al. Pain relief produces negative reinforcement through activation of mesolimbic reward-valuation circuitry. Proceedings of the National Academy of Sciences of the United States of America 109, 20709–20713 (2012). 10.1073/pnas.1214605109

11 Johansen, J. P. & Fields, H. L. Glutamatergic activation of anterior cingulate cortex produces an aversive teaching signal. Nature neuroscience 7, 398–403 (2004). 10.1038/nn1207

12 Zhang, Q. et al. Chronic pain induces generalized enhancement of aversion. eLife 6 (2017). 10.7554/eLife.25302

13 Zhou, H. et al. Ketamine reduces aversion in rodent pain models by suppressing hyperactivity of the anterior cingulate cortex. Nat Commun 9, 3751 (2018). 10.1038/s41467-018-06295-x

14 Hutchison, W. D., Davis, K. D., Lozano, A. M., Tasker, R. R. & Dostrovsky, J. O. Pain-related neurons in the human cingulate cortex. Nature neuroscience 2, 403–405 (1999). 10.1038/8065

15 Iwata, K. et al. Anterior cingulate cortical neuronal activity during perception of noxious thermal stimuli in monkeys. J Neurophysiol 94, 1980–1991 (2005). 10.1152/jn.00190.2005

16 Yamamura, H. et al. Morphological and electrophysiological properties of ACCx nociceptive neurons in rats. Brain research 735, 83–92 (1996).

17 Kung, J. C., Su, N. M., Fan, R. J., Chai, S. C. & Shyu, B. C. Contribution of the anterior cingulate cortex to laser-pain conditioning in rats. Brain research 970, 58–72 (2003).

18 Kuo, C. C. & Yen, C. T. Comparison of anterior cingulate and primary somatosensory neuronal responses to noxious laser-heat stimuli in conscious, behaving rats. J Neurophysiol 94, 1825–1836 (2005). 10.1152/jn.00294.2005

19 Sikes, R. W. & Vogt, B. A. Nociceptive neurons in area 24 of rabbit cingulate cortex. J Neurophysiol 68, 1720–1732 (1992).

20 Zhang, Y. et al. Ensemble encoding of nociceptive stimulus intensity in the rat medial and lateral pain systems. Molecular pain 7, 64 (2011). 10.1186/1744-8069-7-64

21 Hu, S., Zhang, Q., Wang, J. & Chen, Z. Real-time particle filtering and smoothing algorithms for detecting abrupt changes in neural ensemble spike activity. J Neurophysiol 119, 1394–1410 (2018). 10.1152/jn.00684.2017

22 Chen, Z., Zhang, Q., Tong, A. P., Manders, T. R. & Wang, J. Deciphering neuronal population codes for acute thermal pain. Journal of neural engineering 14, 036023 (2017). 10.1088/1741-2552/aa644d

23 Singh, A. et al. Mapping Cortical Integration of Sensory and Affective Pain Pathways. Curr Biol 30, 1703–1715 e1705 (2020). 10.1016/j.cub.2020.02.091

24 Sun, G. et al. Closed-loop stimulation using a multiregion brain-machine interface has analgesic effects in rodents. Sci Transl Med 14, eabm5868 (2022). 10.1126/scitranslmed.abm5868

25 Zhang, Q. et al. A prototype closed-loop brain-machine interface for the study and treatment of pain. Nat Biomed Eng 7, 533–545 (2023). 10.1038/s41551-021-00736-7

26 Qu, C. et al. Lesion of the rostral anterior cingulate cortex eliminates the aversiveness of spontaneous neuropathic pain following partial or complete axotomy. Pain 152, 1641–1648 (2011). 10.1016/j.pain.2011.03.002

27 Gao, Y. J., Ren, W. H., Zhang, Y. Q. & Zhao, Z. Q. Contributions of the anterior cingulate cortex and amygdala to pain- and fear-conditioned place avoidance in rats. Pain 110, 343–353 (2004). 10.1016/j.pain.2004.04.030

28 Li, A. et al. Disrupted population coding in the prefrontal cortex underlies pain aversion. Cell Rep 37, 109978 (2021). 10.1016/j.celrep.2021.109978

29 Ji, G. & Neugebauer, V. Pain-related deactivation of medial prefrontal cortical neurons involves mGluR1 and GABA(A) receptors. J Neurophysiol 106, 2642–2652 (2011). 10.1152/jn.00461.2011

30 Kelly, C. J., Huang, M., Meltzer, H. & Martina, M. Reduced Glutamatergic Currents and Dendritic Branching of Layer 5 Pyramidal Cells Contribute to Medial Prefrontal Cortex Deactivation in a Rat Model of Neuropathic Pain. Frontiers in cellular neuroscience 10, 133 (2016). 10.3389/fncel.2016.00133

31 Radzicki, D., Pollema-Mays, S. L., Sanz-Clemente, A. & Martina, M. Loss of M1 Receptor Dependent Cholinergic Excitation Contributes to mPFC Deactivation in Neuropathic Pain. The Journal of neuroscience : the official journal of the Society for Neuroscience 37, 2292–2304 (2017). 10.1523/JNEUROSCI.1553-16.2017

32 Zhang, Z. et al. Role of Prelimbic GABAergic Circuits in Sensory and Emotional Aspects of Neuropathic Pain. Cell Rep 12, 752–759 (2015). 10.1016/j.celrep.2015.07.001

33 Cheriyan, J. & Sheets, P. L. Altered Excitability and Local Connectivity of mPFC-PAG Neurons in a Mouse Model of Neuropathic Pain. The Journal of neuroscience : the official journal of the Society for Neuroscience 38, 4829–4839 (2018). 10.1523/JNEUROSCI.2731-17.2018

34 Huang, J. et al. A neuronal circuit for activating descending modulation of neuropathic pain. Nature neuroscience 22, 1659–1668 (2019). 10.1038/s41593-019-0481-5

35 Ji, G. & Neugebauer, V. CB1 augments mGluR5 function in medial prefrontal cortical neurons to inhibit amygdala hyperactivity in an arthritis pain model. Eur J Neurosci 39, 455–466 (2014). 10.1111/ejn.12432

36 Lee, M. et al. Activation of corticostriatal circuitry relieves chronic neuropathic pain. The Journal of neuroscience : the official journal of the Society for Neuroscience 35, 5247–5259 (2015). 10.1523/JNEUROSCI.3494-14.2015

37 Martinez, E. et al. Corticostriatal Regulation of Acute Pain. Frontiers in cellular neuroscience 11, 146 (2017). 10.3389/fncel.2017.00146

38 Zhou, H. et al. Inhibition of the Prefrontal Projection to the Nucleus Accumbens Enhances Pain Sensitivity and Affect. Frontiers in cellular neuroscience 12, 240 (2018). 10.3389/fncel.2018.00240

39 Apkarian, A. V. et al. Chronic back pain is associated with decreased prefrontal and thalamic gray matter density. The Journal of neuroscience : the official journal of the Society for Neuroscience 24, 10410–10415 (2004). 10.1523/JNEUROSCI.2541-04.2004

40 Moayedi, M. et al. Contribution of chronic pain and neuroticism to abnormal forebrain gray matter in patients with temporomandibular disorder. NeuroImage 55, 277–286 (2011). 10.1016/j.neuroimage.2010.12.013

41 Geha, P. Y. et al. The brain in chronic CRPS pain: abnormal gray-white matter interactions in emotional and autonomic regions. Neuron 60, 570–581 (2008). 10.1016/j.neuron.2008.08.022

42 Dale, J. et al. Scaling Up Cortical Control Inhibits Pain. Cell Rep 23, 1301–1313 (2018). 10.1016/j.celrep.2018.03.139

43 Sesack, S. R., Deutch, A. Y., Roth, R. H. & Bunney, B. S. Topographical organization of the efferent projections of the medial prefrontal cortex in the rat: an anterograde tract-tracing study with Phaseolus vulgaris leucoagglutinin. The Journal of comparative neurology 290, 213–242 (1989). 10.1002/cne.902900205

44 Liu, Y. et al. Oxytocin promotes prefrontal population activity via the PVN-PFC pathway to regulate pain. Neuron 111, 1795–1811 e1797 (2023). 10.1016/j.neuron.2023.03.014

45 Friedrich, J., Zhou, P. & Paninski, L. Fast online deconvolution of calcium imaging data. PLoS Comput Biol 13, e1005423 (2017). 10.1371/journal.pcbi.1005423

46 Li, X. Y. et al. Alleviating neuropathic pain hypersensitivity by inhibiting PKMzeta in the anterior cingulate cortex. Science 330, 1400–1404 (2010). 10.1126/science.1191792

47 De Felice, M. et al. Capturing the aversive state of cephalic pain preclinically. Annals of neurology (2013). 10.1002/ana.23922

48 King, T. et al. Unmasking the tonic-aversive state in neuropathic pain. Nature neuroscience 12, 1364–1366 (2009). 10.1038/nn.2407

49 Le, A. M., Lee, M., Su, C., Zou, A. & Wang, J. AMPAkines have novel analgesic properties in rat models of persistent neuropathic and inflammatory pain. Anesthesiology 121, 1080–1090 (2014). 10.1097/ALN.0000000000000351

50 Sesack, S. R. & Pickel, V. M. Prefrontal cortical efferents in the rat synapse on unlabeled neuronal targets of catecholamine terminals in the nucleus accumbens septi and on dopamine neurons in the ventral tegmental area. The Journal of comparative neurology 320, 145–160 (1992). 10.1002/cne.903200202

51 Sotres-Bayon, F., Sierra-Mercado, D., Pardilla-Delgado, E. & Quirk, G. J. Gating of fear in prelimbic cortex by hippocampal and amygdala inputs. Neuron 76, 804–812 (2012). 10.1016/j.neuron.2012.09.028

52 Jhang, J. et al. Anterior cingulate cortex and its input to the basolateral amygdala control innate fear response. Nat Commun 9, 2744 (2018). 10.1038/s41467-018-05090-y

53 Totah, N. K., Jackson, M. E. & Moghaddam, B. Preparatory attention relies on dynamic interactions between prelimbic cortex and anterior cingulate cortex. Cereb Cortex 23, 729–738 (2013). 10.1093/cercor/bhs057

54 Hardy, S. G. & Haigler, H. J. Prefrontal influences upon the midbrain: a possible route for pain modulation. Brain research 339, 285–293 (1985).

55 Qi, C. et al. Anterior cingulate cortex parvalbumin and somatostatin interneurons shape social behavior in male mice. Nat Commun 16, 4156 (2025). 10.1038/s41467-025-59473-z

56 Koga, K. et al. Coexistence of two forms of LTP in ACC provides a synaptic mechanism for the interactions between anxiety and chronic pain. Neuron 85, 377–389 (2015). 10.1016/j.neuron.2014.12.021

57 Xu, H. et al. Presynaptic and postsynaptic amplifications of neuropathic pain in the anterior cingulate cortex. The Journal of neuroscience : the official journal of the Society for Neuroscience 28, 7445–7453 (2008). 10.1523/JNEUROSCI.1812-08.2008

58 Tan, L. L. et al. Gamma oscillations in somatosensory cortex recruit prefrontal and descending serotonergic pathways in aversion and nociception. Nat Commun 10, 983 (2019). 10.1038/s41467-019-08873-z

59 Tan, L. L. et al. A pathway from midcingulate cortex to posterior insula gates nociceptive hypersensitivity. Nature neuroscience 20, 1591–1601 (2017). 10.1038/nn.4645

60. Alonso-Matielo, H., et al. Inhibitory insula-ACC projections modulate affective but not sensory aspects of neuropathic pain. Mol Brain 16, 64 (2023). 10.1186/s13041-023-01052-8

61 Chen, Z. S. Hierarchical predictive coding in distributed pain circuits. Front Neural Circuits 17, 1073537 (2023). 10.3389/fncir.2023.1073537

62 Zhou, P. et al. Efficient and accurate extraction of in vivo calcium signals from microendoscopic video data. eLife 7 (2018). 10.7554/eLife.28728

63 Pnevmatikakis, E. A. et al. Simultaneous Denoising, Deconvolution, and Demixing of Calcium Imaging Data. Neuron 89, 285–299 (2016). 10.1016/j.neuron.2015.11.037

64 Swietek, B., Gupta, A., Proddutur, A. & Santhakumar, V. Immunostaining of Biocytin-filled and Processed Sections for Neurochemical Markers. J Vis Exp (2016). 10.3791/54880

65 Goffer, Y. et al. Calcium-permeable AMPA receptors in the nucleus accumbens regulate depression-like behaviors in the chronic neuropathic pain state. The Journal of neuroscience : the official journal of the Society for Neuroscience 33, 19034–19044 (2013). 10.1523/JNEUROSCI.2454-13.2013

